# Open-source tag-free monitoring of individual birds using automated weighing and deep-learning recognition

**DOI:** 10.64898/2026.08.17.745158

**Authors:** Jinook Oh, Marisa Hoeschele

## Abstract

Effective animal monitoring is essential for assessing health, behavior, and environmental interactions, particularly in research and welfare contexts. This study presents a low-cost, open-source system designed for non-invasive monitoring of budgerigars (*Melopsittacus undulatus*), a small parrot species frequently used in animal behavior research. The system integrates a perch-based scale for voluntary weight measurement, a temperature sensor, and a camera for image capture, all controlled by a Raspberry Pi. By leveraging fine-tuned neural networks, the system achieves automated individual recognition with high accuracy, eliminating the need for invasive tagging methods. The modular design ensures accessibility, scalability, and minimal disturbance to the animals, while the accompanying software streamlines data collection, processing including labeling, and visualization. This approach provides a comprehensive solution for continuous monitoring, offering valuable insights for research and husbandry while prioritizing animal welfare.

## 1 Introduction

Monitoring and observing animals is essential not only for assessing their health but also for generating insights that can lead to new research questions [1, 2, 3]. Effective animal monitoring provides critical data on individual health, behavior, and environmental interactions, forming the foundation for both scientific studies and welfare practices. Among the various health indicators, body weight is one of the most crucial metrics [4, 5, 6, 7]. Tracking the temporal fluctuations in an animal’s weight can reveal valuable information about its health state, nutritional status, and overall well-being. Because small psittacines like budgerigars (*Melopsittacus undulatus*) have limited body-mass reserves and high metabolic rates, even minor fluctuations in weight can indicate emerging illness. Several studies have emphasized the importance of regular weighing and body-condition assessment in small bird species to monitor health status [5, 8]. Additionally, monitoring environmental factors such as temperature is vital for maintaining optimal conditions, particularly for species such as budgerigars (*Melopsittacus undulatus*), which are sensitive to temperature fluctuations in captivity [9].

Traditionally, weighing animals involves manually capturing the animal, placing it in a small cage, and measuring the combined weight of the cage and the animal using a commercial scale. While this method is straightforward, it comes with significant drawbacks. The physical capture of animals can be highly stressful for them, potentially altering their behavior and physiological state [10, 11]. Additionally, this process requires trained personnel, and considerable human effort, and time, making it impractical for frequent or large-scale monitoring.

Another challenge in weight monitoring is the need to identify which individual is being weighed. Various tagging methods, such as coloring, RFID chips, QR codes, or other artificial markers, are commonly used for individual identification [12, 13, 14, 15]. However, these methods are not always feasible depending on species traits or experimental limitations. For instance, budgerigars which weigh only 30–40 grams, are too small to carry most artificial tags. Even lightweight RFID tags, such as a 1*cm* long passive RFID chip, present significant challenges. These tags often require the antenna to be in very close proximity—1–3*cm* depending on the alignment of the antenna and the tag—for successful reading, which can be impractical in many scenarios. Additionally, securely attaching the tag without causing discomfort or harm to the bird is difficult. The tag must be highly durable (to withstand the bird’s beak) and lightweight to avoid interfering with the bird’s natural behavior. Similarly, artificial color tags come with their own set of challenges. While they do not require specialized reading equipment, the natural color patterns in budgerigars are important and artificial color marking can alter how the bird is perceived by its peers, potentially disrupting social interactions and colony dynamics [16, 17]. This behavioral impact makes color tagging less suitable for species such as budgerigars, where maintaining natural social behavior is critical when the research is related to its social behavioral aspects. Manual identification of individuals based on their natural appearance also poses challenges, particularly in group settings where animals frequently interact and move.

Considering these challenges, a system that enables voluntary and automatic weighing, combined with individual identification based on natural appearance, would be highly beneficial. Such a system would not only reduce stress for the animals but also eliminate the need for invasive tagging or manual handling. Moreover, continuous monitoring of weight, paired with automated identification, would provide a wealth of data for both health assessments and behavioral studies. The integration of image data into this process would further enhance its utility, enabling researchers to use them in different projects as a base dataset. Additionally, these features are particularly valuable for longitudinal studies, where tracking the health and behavior of individuals over extended periods can yield critical insights into life history traits, aging, and the effects of environmental changes.

In this study, we propose a novel device and pipeline designed to address these needs. Our system combines a perch-based scale for voluntary weighing, temperature monitoring, and image capture for the neural network-based identification. While the system is specifically designed for budgerigars, its modular and adaptable nature makes it suitable for broader applications in monitoring other small animals. This approach ensures non-invasive, automated, and continuous monitoring, while providing valuable data for research and husbandry. By leveraging open-source software and affordable hardware, the system is accessible and scalable, making it a versatile tool for a wide range of applications in animal monitoring, welfare, and conservation.

## 2 Materials and Methods

### 2.1 Animal species

The monitoring solution in this study was designed specificially for budgerigars (*Melop-sittacus undulatus*). The budgerigar, commonly known as the budgie, is a small parrot species native to Australia. Their small size, ease of care, and adaptability to captive environments have made them a preferred species for studies in avian biology, animal cognition and neuroethology [18, 19, 20, 21, 22]. Visual identification of individual budgerigars is an interesting recognition problem because human experts appear to rely on holistic, face-like visual processing mechanisms rather than isolated visual features. This is supported by findings that inversion of budgerigar images—turning them upside down—significantly impairs experts’ ability to recognize individuals, demonstrating a configural processing effect similar to the face inversion effect in humans [23]. Comparable inversion effects have also been observed in other contexts of animal individual recognition, including expert identification of dogs and conspecific face discrimination in sheep [24, 25]. In addition, budgerigars offer a practical advantage as an experimental model because their small body size allows large numbers of individuals to be housed and observed simultaneously under laboratory conditions. Although budgerigars often display subtle yet individually distinctive plumage patterns, expert recognition relies on holistic processing of these patterns and their spatial configurations rather than isolated features. These characteristics make budgerigars an attractive model for developing and evaluating machine-learning approaches that capture complex spatial relationships for individual identification. Consequently, they provide an ideal species for developing and validating monitoring systems such as the one presented in this study.

### 2.2 Tools for development

All the programs for this study were developed in Python (version 3.13, retrieved fromhttps://www.python.org) with several essential packages; OpenCV (version 4.11, retrieved fromhttps://opencv.org), wxPython (version 4.2, retrieved fromhttps://wxpython.org), PyTorch (version 2.6, retrieved fromhttps://pytorch.org), NumPy (version 2.2, retrieved fromhttps://numpy.org), RPi.GPIO (version 0.7, retrieved fromhttps://pypi.org/project/RPi.GPIO), picamera (version 1.1, retrieved from https://pypi.org/project/picamera) and HX711 (version 1.0, retrieved fromhttps://github.com/gandalf15/HX711). 3D printed parts in hardware were designed in Blender (4.2, retrieved fromhttps://www.blender.org) and printed using a FDM (Fused Deposition Modeling) 3D printer (Ultimaker, Netherlands). A developed software for monitoring functionalities were installed and run on a single-board computer (hereafter SBC; Raspberry Pi 4b 4GB RAM, Raspberry Pi Foundation, United Kingdom). A load cell (Dioche, China) and HX711 (24-Bit Analog-to-Digital converter for weigh scales; Hailege, China) were used together to acquire a budgerigar’s weight. A camera (C920, Logitech, Switzerland) was attached to the SBC for the image acquisition.

### 2.3 Hardware

The monitoring unit was designed to be compact, functional, and easy to assemble using readily available components. The hardware consists of a custom-built scale, a temperature sensor, and a camera, all connected to an SBC for data collection and processing.

The scale was constructed using a combination of 3D-printed and hand-carved wooden components. Two 3D-printed pieces were designed to securely hold the load cell in place and provide structural support for the perch. The wooden perch, which was hand-carved to ensure a natural and comfortable surface for the budgerigars, while not allowing two budgerigars at the same time with its short length; 7*cm*, was mounted on the load cell. The load cell (100 g capacity) was chosen for its sensitivity and ability to measure small weight changes accurately. The load cell was connected to an HX711 amplifier, which converts the analog signals from the load cell into digital signals for processing. The HX711 was then connected to the SBC via the GPIO pins, specifically using the 3.3V power pin, ground (GND), and two data pins (DT and SCK) for communication. As budgerigars are a social species, a regular perch was placed in close proximity to the perch scale to ensure that the budgerigar on the scale does not feel isolated.

To monitor the environmental temperature, a DS18B20 digital temperature sensor was integrated into the system. This sensor was connected to the GPIO pins of the SBC and configured to measure the temperature at regular intervals. In a file, /boot/firmware/-config.txt, the following line was added, `dtoverlay=w1-gpio,gpiopin=16,pullup=on`, as pin-16 was used for the temperature sensor in this study. The DS18B20 was selected for its accuracy, ease of use, and compatibility with the SBC. It provides reliable temperature readings, which are essential for maintaining the indoor environment of the aviary. Although wild budgerigars originate from semi-arid Australian habitats with frequent temperatures well above typical indoor levels, captive budgerigars used in laboratory and husbandry settings are less tolerant and routinely housed at room temperatures of *≈* 20-25 °C [26, 9].

A Logitech C920 HD Pro webcam was used to capture images of the budgerigars during weight measurements. The camera was connected to the SBC via a USB 3.0 port and positioned on top of the aviary, approximately 50 *cm* above the perch, with an unobstructed view to ensure clear imaging of birds on the perch and sufficient image resolution for the subsequent visual identification task.

The Raspberry Pi was chosen for its versatility, affordability, and compatibility with the various sensors and peripherals. It was responsible for scheduling of signal reading, processing data from the 1) load cell, 2) temperature sensor and 3) camera, and storing the collected data, via a Python script.

All components were carefully assembled and mounted on the aviary wall and ceiling to ensure stability and functionality. The design was optimized to avoid interfering with the birds’ natural activities while also protecting the wires and hardware from potential damage caused by the birds (See Figure 1).

**Figure 1:**
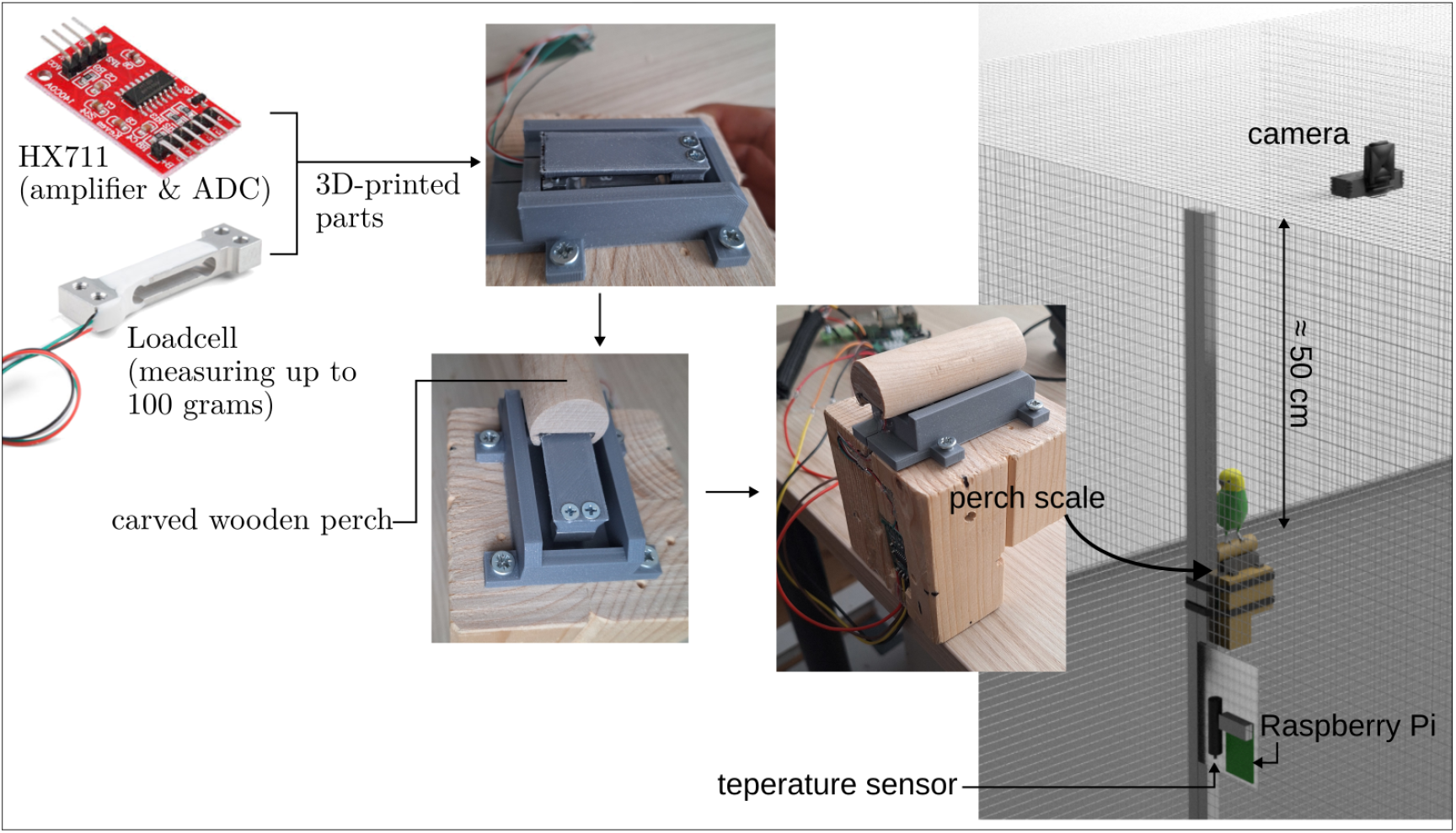
Monitoring unit.

### 2.4 Software

All the code developed for this study was written in Python and is open-source. The code and its data are made publicly available on Github (https://github.com/jinook0707/code4Ethology) and Zenodo (https://doi.org/10.5281/zenodo.19351938) to ensure reproducibility and accessibility.

The software was designed to automate the monitoring of temperature, weight, and image capture, integrating seamlessly with the hardware components described earlier. Furthermore, the software includes tools for efficiently labeling the captured budgerigar images and for training (fine-tuning) a backbone neural network. This allows the creation of a model specialized in the identification of a specific set of individual birds, enabling automated and accurate recognition based on the collected data.

#### 2.4.1 Code on SBC; monitorAviary.py

The script, monitorAviary.py, was configured to run automatically in the background upon the SBC boots up by registering it as a systemd service, defined in `/etc/systemd/system/mAviary.service`, which specifies settings such as the working directory, execution command, restart policy, and standard output handling. The contents of `mAviary.service` is written in Appendix A, which ensures that `monitorAviary. py` operates reliably, after system reboots due to power interruptions or other causes.

The script, `monitorAviary.py`, includes configurable parameters, such as reading intervals, GPIO pins and calibration factors for accessing data from the temperature and the custom scale. These parameters should be adjusted whenever the hardware setup changes — for example, when GPIO pins are reassigned, sensors are replaced, or recalibration is required. The script periodically reads data from the DS18B20 temperature sensor at user-defined intervals. Each temperature reading is recorded in a daily output file, named based on the current date (e.g., `temperature_YYYY-MM-DD.log`). Each entry in the file includes a timestamp and the corresponding temperature value. This allows for continuous tracking of environmental conditions in the aviary, ensuring that the temperature remains within the optimal range for the budgerigars. Furthermore, the logged temperature data can be used to correlate ambient conditions with observed behaviors or physiological measures, providing insights into how environmental temperature influences the wellbeing of the budgerigars.

The weight monitoring process is designed to detect when a bird is sitting steadily on the perch and to record its weight along with a corresponding image for identification. The logical steps for weight monitoring are outlined in Algorithm 1. The process is as follows:

##### Algorithm 1

Weight reading

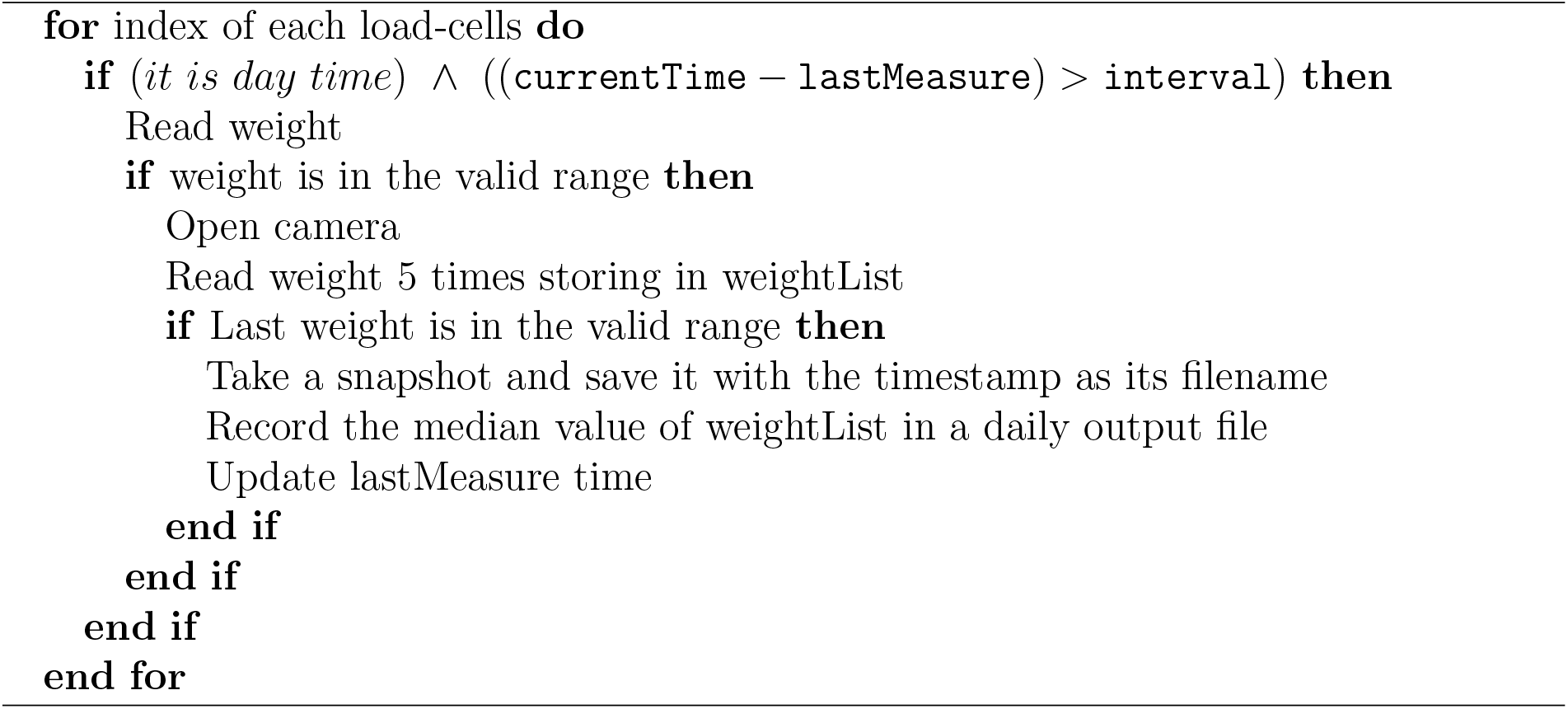

#### 2.4.2 Namer

`Namer` is a newly developed GUI application created for this project. It utilizes wxPython, the Python implementation of the cross-platform wxWidgets C++ library. The interface of `Namer` is shown in Figure 2. The user can open a folder, which has the snapshot image (1920×1080) files to identify and corresponding log files. When the app opens the folder, it loads the images and crops the image around the perch if it’s not cropped yet, i.e.: the image size is 1920×1080. The rect for cropping is set in `settings.json` file, which has user defined parameters such as `modelName, lossFunc, wghtFN, xywh4crop, budgieNames`.

**Figure 2:**
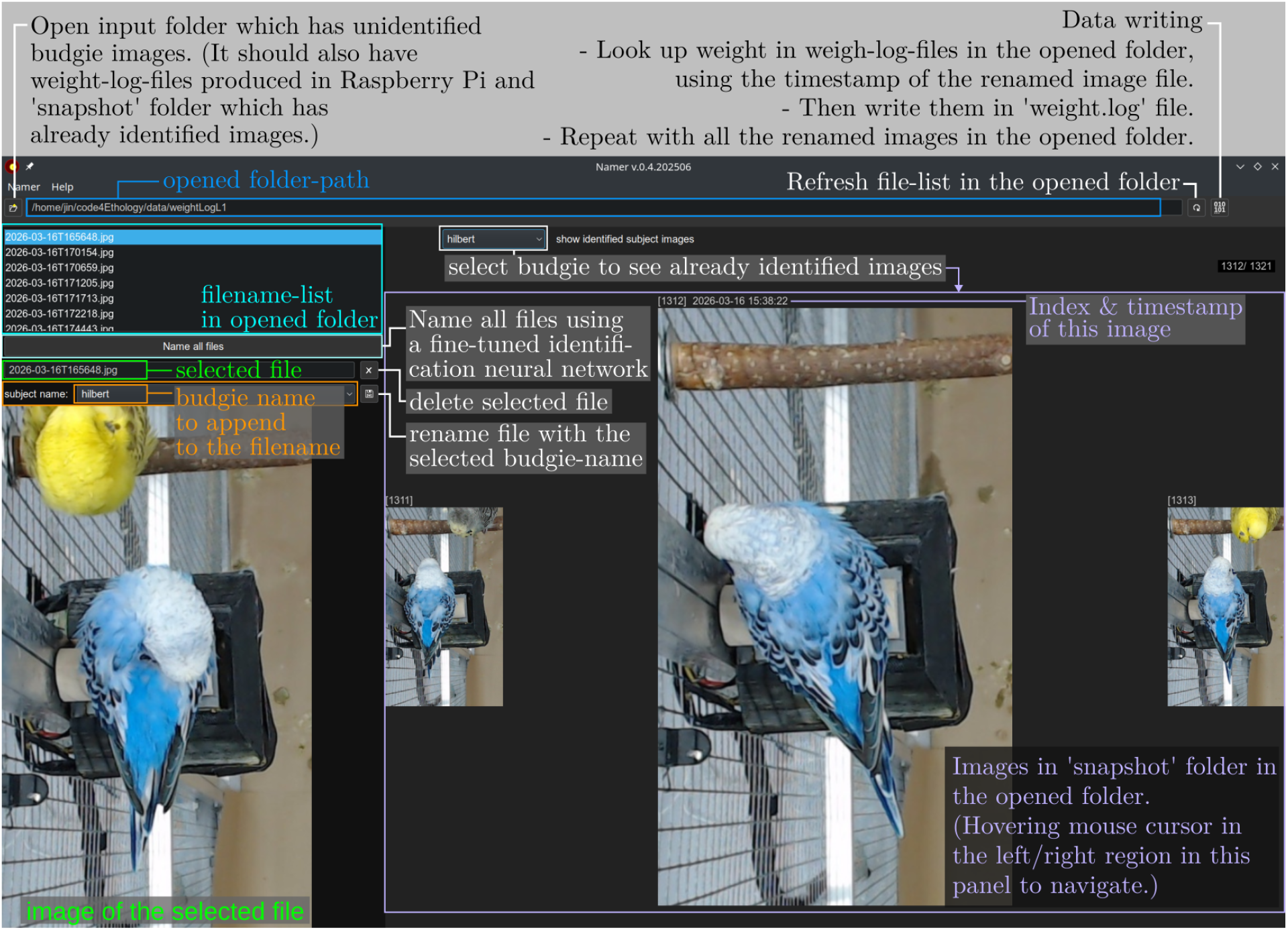
GUI user application; Namer.

##### Initial labeling

In the initial manual labeling phase of `Namer`, the user is responsible for reviewing and labeling the captured images of birds. If no bird is present on the perch in an image, the user can press the `Delete` button to remove the corresponding image file. For images where a bird is present, the user selects the bird’s name from the ‘subject name’ drop-down list of choices. These choices are defined in the `budgieNames` field of the `settings.json` configuration file, allowing for easy customization to match the specific individuals being monitored.

If the identity of the bird in an image is unclear, the user can use the right-side panel to select a bird name and view all previously labeled images of individuals from the `snapshots` folder in the input directory. This feature provides a visual reference to assist in resolving ambiguities during the labeling process. After the correct name is chosen for the bird, the user clicks the button with the save icon, which renames the original file by appending the bird name at the end, resulting in a ‘[timestamp] [subject-name].jpg’ filename.

Once all images have been manually labeled, the user can press the data saving button. This action searches for the timestamps of each snapshot, matches them with the corresponding weight data in the weight log file, and appends a line to the final data file, `weight.log`. Each line in the file consists of the timestamp, the bird’s name, and the associated weight data. As part of this process, all image files are moved into the ‘snapshot’ folder, and the raw weight log files are relocated to the ‘rawLogFiles’ folder, ensuring the input directory is organized and ready for future data processing. Then, the user can proceed to the next phase of the workflow, where the labeled data can be used to fine-tune neural network models for automated identification.

##### Neural Network

Nine different image classification models, often used for image identification tasks, were downloaded and tested using TIMM (pyTorch IMage Models), which is a comprehensive library of pre-trained image models, layers, optimizers, utilities, etc. The models tested were EfficientNet b0, MobileNet v3, VisionTransformer (ViT) Tin y, EfficientNet v2 b0, EfficientNet v2 b3, VisionTransformer (ViT) Small, Swin Transformer Tiny, ConvNeXT Tiny, and RegNetY-6.4GF, listed in the order of increasing parameter size.

A total of 539 snapshot images of budgerigars, voluntarily sitting on the scale, were collected over a period of approximately four months. The number of images collected per individual bird varied significantly, with counts of 4, 4, 6, 15, 20, 33, 41, 52, 64, 105, and 195 images for each bird. To create a balanced training dataset, 100 training images per class (1,200 images in total for 11 individual budgerigars and an empty perch) were generated. For birds with fewer than 100 images, the training pipeline automatically applied data augmentation, consisting of random selections from the original set followed by random flipping, rotation, and translation. For birds with more than 100 images, a random subset of 100 images was selected from the original set. Example images are shown in Figure 3-(a).

**Figure 3:**
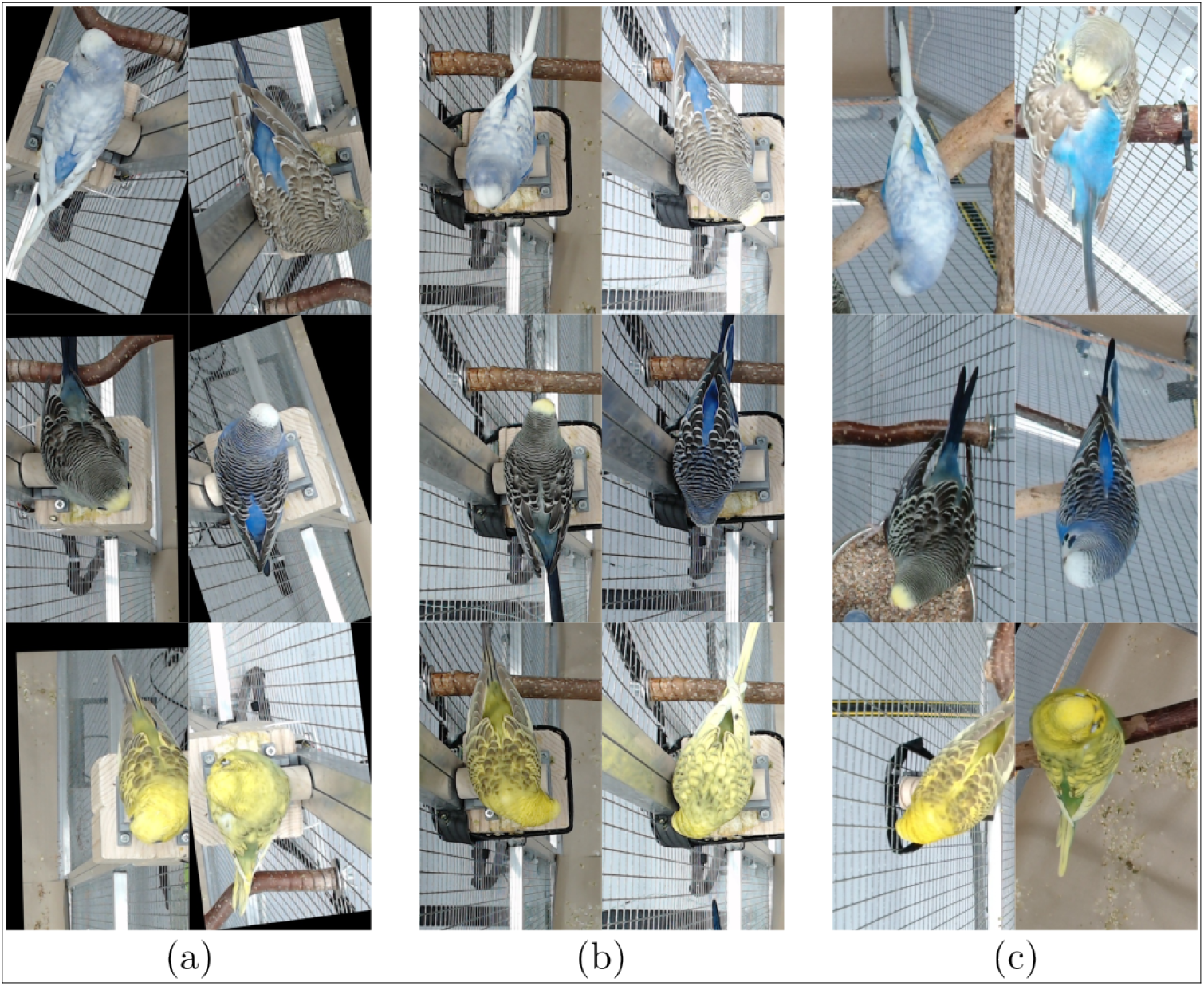
Example images from (a) dataset for training (fine-tuning), (b) dataset 1 and (c) dataset 2 for testing.

Using the generated training dataset, each model was first trained using the softmax classification approach, which is the standard method for classification tasks. Subsequently, the models were also trained using metric learning to evaluate whether this approach could better generalize to new datasets, particularly those with different backgrounds from the training set.

To test the trained models, two distinct datasets were used:

Dataset 1: This dataset consisted of 122 images of budgerigars sitting on the same scale used during training (see Figure 3-(b)).

Dataset 2: This dataset consisted of 330 images (30 images per budgerigar), which included a mix of budgerigars sitting on a different scale (with a different location and design) and on various perches in different locations within the aviary (see Figure 3-(c)).

The images of budgerigars on perches were extracted from videos collected for another study, which utilized an array of SBCs with 18 cameras mounted above the aviary. To facilitate the extraction of budgerigar images from videos, a custom GUI tool, `sampleImgExtract`. py, was developed. It allows users to define a rotated rectangular region of interest with fully adjustable width, height, and angle via keyboard input. The extracted images were then used to create Dataset 2.

The performance of each model, trained using either the softmax or metric learning method, was evaluated on both datasets. The performance measure was the percentage of correct inferences made by the model.

##### AI assisted labeling

After training, several parameters `in settings.json` were specified, including `modelName, lossFunc` (softmax or metric), and `wghtFN` (name of the weight file). If the file `weightFiles_[lossFunc]/[wghtFN]` exists, the Namer program uses the specified classifier to assist in identifying the budgerigar’s name.

When a budgerigar image is selected, the program reorders the ‘subject name’ drop-down list based on the classifier’s predicted probabilities, placing the most likely names at the top. If the classification is correct, the user can efficiently label the image by clicking the save (file-renaming) button. Furthermore, if the classifier’s performance is sufficiently reliable, the user may utilize the ‘Name all files button (see Figure 2)’ to rename all files automatically based on the classifier’s predictions, bypassing the need for manual input.

##### Data visualization

For specialized visualization of the collected temperature and weight data, a GUI application called `Visualizer` was used. Although `Visualizer` was originally developed for a previous study [27], its graph-drawing functionality was modularized into a single function or class, allowing additional features to be implemented with minimal effort. This flexibility enabled the creation of customized visualizations tailored to the specific needs of this study and is included in the current codebase.

## 3 Results

The temperature and weight monitoring unit has been active from December 2024 to the current date. Example resultant graphs drawn by the `Visualizer` are shown in Figure 4 and 5.

**Figure 4:**
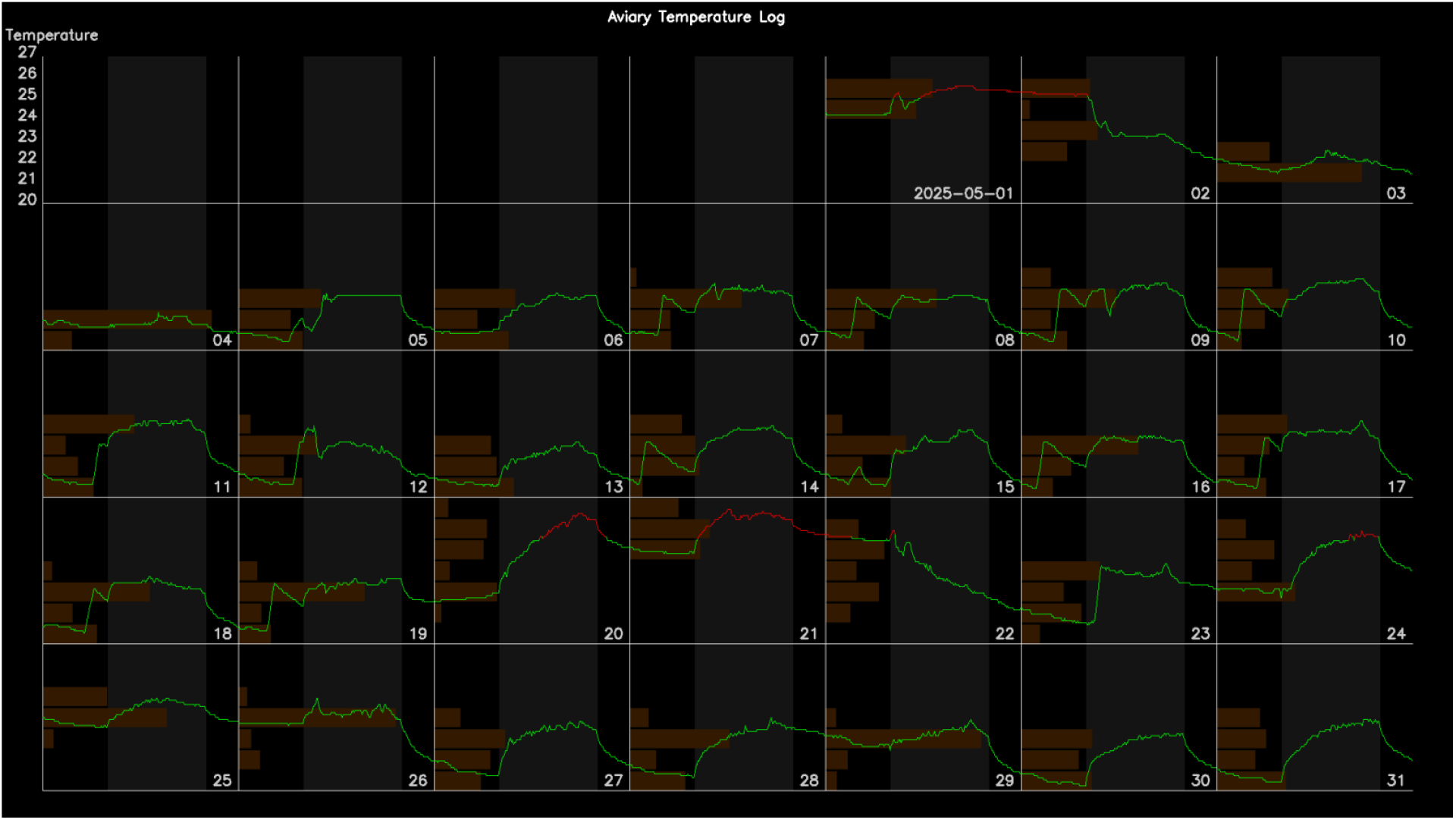
Example temperature graph, compiled from minute-by-minute log data aggregated at 5-minute intervals using median values. Red segments denote the temperature out of the normal range (20–25 °C). Horizontal histograms along each day illustrate the frequency distribution of temperature readings across 1 °C bins, providing an overview of daily temperature stability and variability.

**Figure 5:**
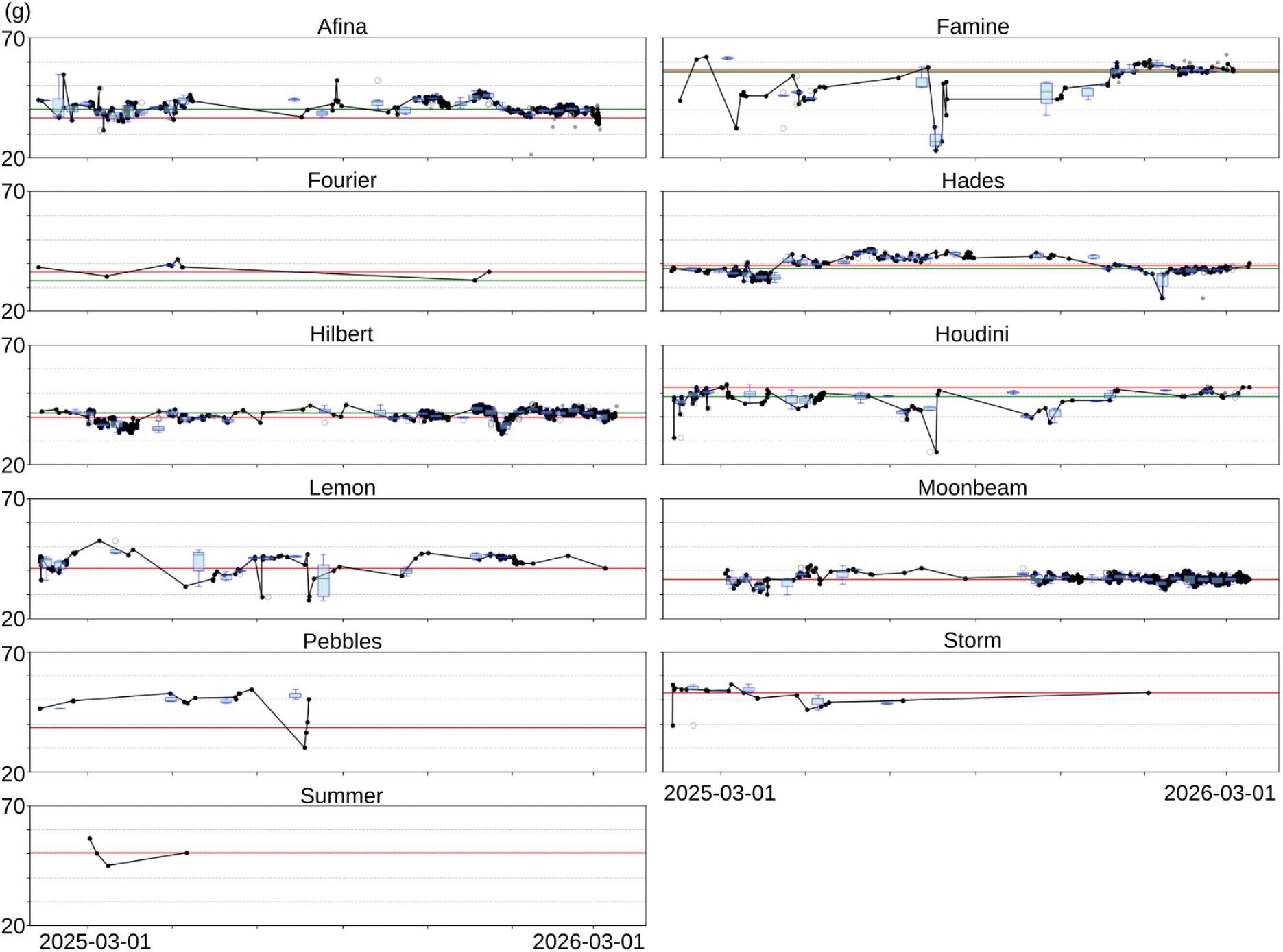
Weight graph over an approx. 1 year period; Each subplot corresponds to one bird, showing all recorded body weights over time (black dots). Outlier values, identified using a rolling z-score threshold (*σ* = 2) within 10-day windows, are shown in grey. Blue boxplots summarize the distribution of body weights within each 10-day interval. The solid green horizontal line spans the entire subplot and represents the median weight of the preceding 10-day period, while the red line indicates the median of the most recent 10-day period, enabling a direct visual comparison of recent weight shifts.

### 3.1 Comparison of neural models for individual identification

Figure 6 shows the comparison results of each neural model either with softmax or metric learning on both dataset 1 and 2. To assess whether balanced batch sampling improves metric learning performance, an additional test (`metric_D2_b`) was conducted on dataset 2. On dataset 1, many models achieved near-perfect identification accuracy (about 99–100%), indicating that they can identify individual appearances when background conditions are similar to those in the training images. In contrast, performance on dataset 2 diverged sharply. Smaller convolutional models (< ∼ 20 million parameters) trained with a standard softmax classifier showed substantial drops in accuracy (e.g., MobileNet / EfficientNet-B0: 60–75%). When trained with a metric-learning objective, however, the same architectures improved markedly (about 89–96%). Larger models (*>* 20 million parameters; ViT-small, ConvNeXt-tiny, RegNet-Y64) maintained strong performance across both datasets, with ViT-small and ConvNeXt-tiny showing the highest accuracy.

**Figure 6:**
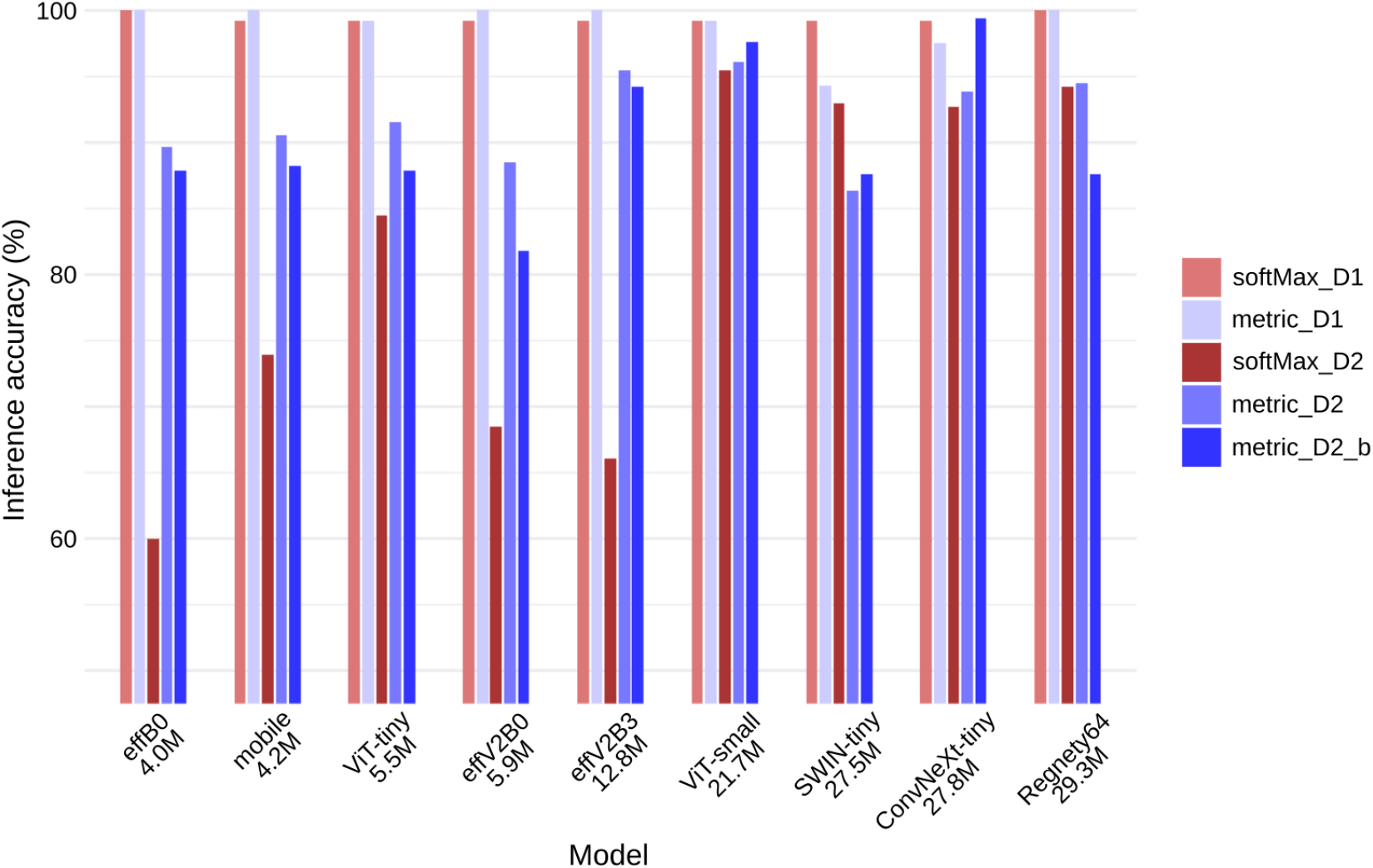
Comparison of neural models on the individual bird identification task * D1 and D2 are the dataset 1 and 2 explained in Section 2.4.2. * `metric_D2_b` is metric learning with balanced batch sampling, tested on D2 * Models are in order of increasing parameter size.

Balanced batch sampling further benefited several larger models (ViT-small, SWINtiny, and ConvNeXt-tiny), with ConvNeXt-tiny showing the most marked improvement. Based on these results, we selected the ConvNeXt-tiny model trained with metric learning and balanced batch sampling for integration into `Namer`. In addition to overall classification accuracy (*≈*99.4%), the performance of the selected model was evaluated using the macro-F1 score. The macro-F1 score was 0.994 and was calculated as the unweighted mean of the class-wise F1 scores across individuals. A confusion matrix (Appendix B) was generated to examine the distribution of misclassifications among individuals. Notably, the ConvNeXt-tiny model retained its high (≈99%) accuracy on over 1000 images when the scale’s color and shape were later modified, demonstrating robustness to such appearance variations.

Incorrect classifications arose from challenging postures — such as when a bird tucked its head beneath its wing — causing its distinctive visual features to become obscured, although such images remained correctly recognizable to the human eye. An illustrative example of such a case is presented in Figure 7. Additional misclassifications occurred when another bird approached in close proximity to the scale.

**Figure 7:**
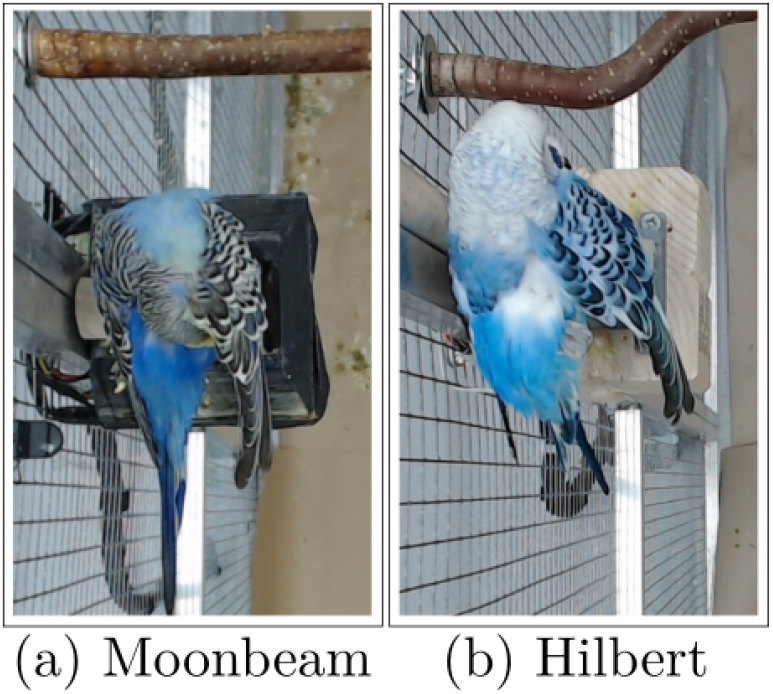
A misclassification case where Moonbeam (a) was falsely predicted as Hilbert; (b) shows a reference image of Hilbert for comparison.

The choice of model depends on the specific goals and setup of each experiment. The model used by Namer is configurable; users can select a different trained model simply by specifying the corresponding weight file. In general, ViT-small or ConvNeXt-tiny are recommended for robust inference, provided that a slightly longer inference time is acceptable. Table 1 reports the mean inference time per image for each model, benchmarked on a Raspberry Pi 5 (8 GB RAM), which is a more recent model than the one deployed in the scale device.

**Table 1:** Mean inference time per image for each model on dataset 2, measured on a Raspberry Pi 5.

| Model | Mean inference time (in seconds) |
| --- | --- |
| efficientnet_b0 | 0.113 |
| mobilenetv3_large_100 | 0.074 |
| vit_tiny_patch16_224 | 0.070 |
| tf_efficientnetv2_b0 | 0.096 |
| tf_efficientnetv2_b3 | 0.263 |
| vit_small_patch16_224 | 0.223 |
| swin_tiny_patch4_window7_224 | 0.297 |
| convnext_tiny | 0.263 |
| regnety_064 | 0.416 |

### 3.2 Reduced time and effort

By integrating the fine-tuned weights – which achieved a classification accuracy exceeding 99% – into `Namer`, the application facilitates an efficient labeling workflow through both individual name suggestions and an autonomous batch-processing feature. This latter functionality allows for the immediate labeling of all images within a target directory in a single operation, leveraging the model’s high predictive performance to bypass manual per-image confirmation. This automation yields a substantial reduction in the temporal workload. Consequently, the user’s role transitions from repetitive manual data entry to high-level verification and occasional oversight.

## 4 Discussion

This study successfully implemented a low-cost, open-source system for non-invasive monitoring of budgerigars, integrating weight measurement, temperature monitoring, and neural network-based individual identification. One of the the key achievements of this work lies in demonstrating that high identification accuracy (*>* 99%) can be achieved using a limited training dataset.

This high identification accuracy with a small sample dataset addresses a critical challenge faced by many biologists: the difficulty of obtaining large, high-quality datasets in real-world research settings. The ability to achieve robust performance with a small dataset is particularly significant for biological and ecological studies, where collecting extensive image data is often impractical due to time, resource, or logistical constraints. By leveraging fine-tuned neural network models and metric learning, it effectively extracts intrinsic visual traits, such as feather patterns and body shape, while minimizing reliance on contextual cues such as background or lighting. This capability not only enhances the system’s generalizability to new conditions but also underscores its potential for broader applications in animal monitoring and welfare research. The results highlight the value of combining high-quality data with advanced machine learning techniques to overcome the limitations of small datasets, enabling accurate and reliable individual identification. One mortality event was recorded in the aviary during the study period. The subject, Afina, was found on the floor on March 6th and was promptly transferred to a veterinary hospital, where he passed away the same day. Notably, caretakers had not observed any significant changes in his behavior or body weight prior to the incident. However, Figure 8 shows a decline in weight during the January–March 2026 period, including four abnormally low weight records — marked as grey dots to denote outlier data points — as well as an episode of sharp weight recovery and drop over the final days. Since avian body weight is subject to natural fluctuation and data collection errors may occur, a detection algorithm relying solely on weight changes risks generating an excessive number of false alarms. Nevertheless, integrating weight monitoring with behavior pattern analyses — such as perch relocation frequency and daily social contact counts — through the aforementioned identification pipeline may offer a more robust approach to predicting health deterioration in birds at an earlier stage.

**Figure 8:**
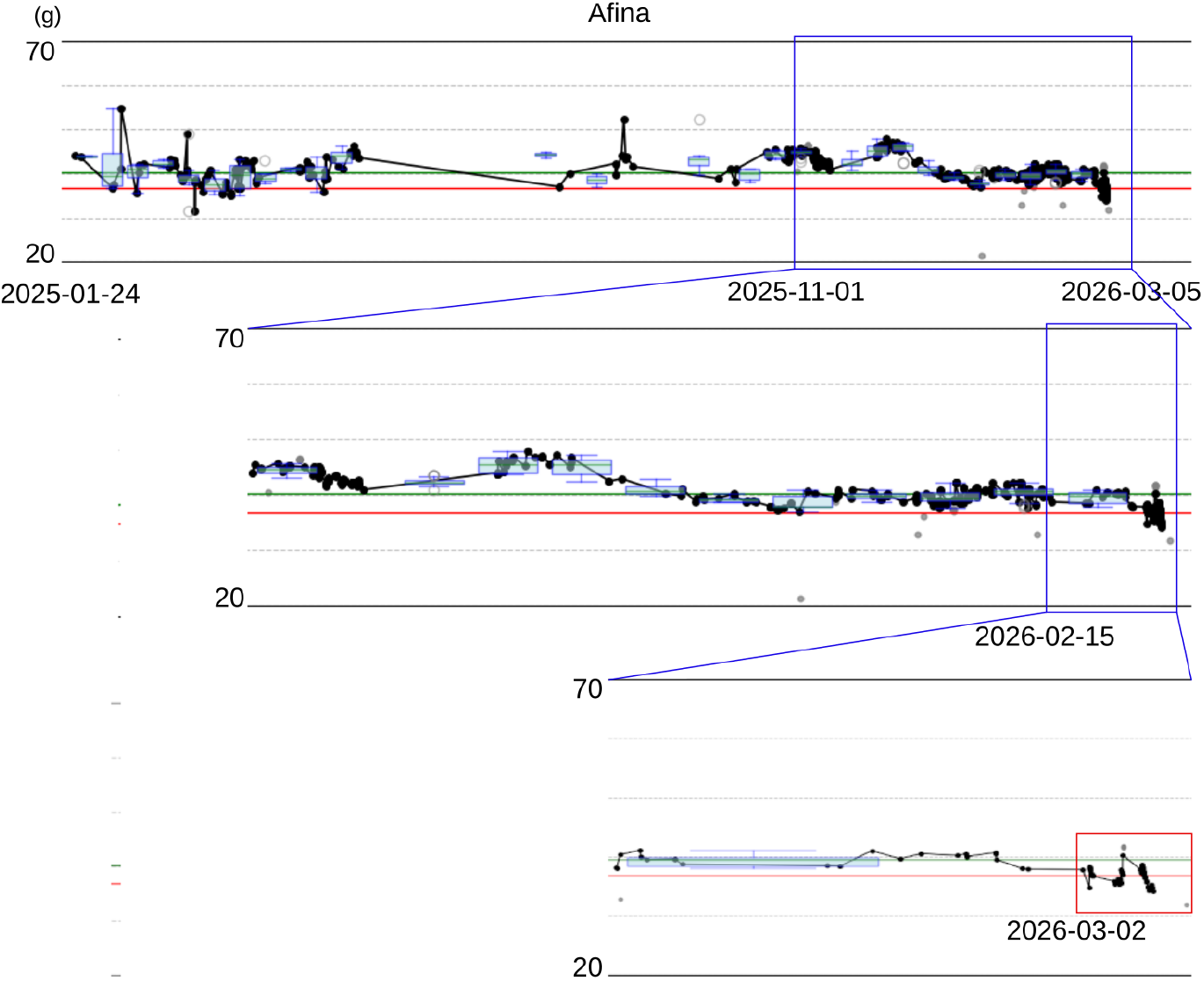
Weight fluctuation of Afina

The system’s affordability and open-source design ensure accessibility for a wide range of researchers. By utilizing widely available components such as the Raspberry Pi, DS18B20 temperature sensor, and Logitech C920 camera, the system is cost-effective and easy to replicate. The perch-based scale provides a natural and comfortable spot for the budgerigars to sit, while its short length ensures that only one budgerigar can perch at a time, enabling reliable weight measurements. Additionally, the user-friendly graphical interface, developed using `wxPython`, simplifies operation, making it particularly suitable for biologists without technical expertise. The `Namer` application allows users to open folders containing snapshot images and log files, automatically crop images when necessary, and label them with minimal effort. Features such as a dropdown menu for selecting bird names and a visual reference panel for previously labeled images simplify the labeling process, making it easy to resolve ambiguities. Automated functionalities, including reordering names or renaming all files based on classifier predictions and matching timestamps with weight data, further streamline the workflow. These thoughtful design elements ensure that the system can be effectively operated by individuals without technical expertise, making it accessible to a wide audience of researchers and enthusiasts.

The system prioritizes animal welfare by encouraging natural behavior through a perch-based scale and non-invasive monitoring methods. The perch-based scale allows birds to weigh themselves voluntarily without external coercion, minimizing stress and ensuring accurate weight measurements. When combined with temperature monitoring, the system provides a comprehensive overview of the aviary’s environmental conditions and the health of its inhabitants. This welfare-oriented design aligns with best practices in research and husbandry, ensuring that the system not only collects valuable data but also safeguards the health and comfort of its subjects.

While designed for budgerigars, the system’s modular and adaptable nature makes it suitable for monitoring other small animals. The captured images, which are well-sighted and high-quality, are ideal for automated identification using advanced neural networks. Recent advancements in machine learning and computer vision have made it possible to identify individuals based on subtle visual traits, such as feather patterns or body shape, without the need for artificial tags. These images can also support studies on social interactions, behavioral patterns, or long-term health monitoring, further expanding the system’s utility.

Considering the performance differences we observed among neural networks, one likely explanation is that softmax classifiers trained on a limited set of images under uniform conditions tend to rely on background or lighting cues linked to each individual. When the background changes, these “shortcut” cues no longer apply, leading to poor generalization. Metric learning, in contrast, trains the network to group images of the same individual close together in a learned feature space while pushing apart different individuals. This tends to emphasize intrinsic visual traits—such as feather patterns, body shape, or markings—rather than contextual information such as background or posture. As a result, metric-learning models are often more robust to background changes and better suited for fine-grained identification tasks with few samples per individual, as shown previously in animal and human re-identification studies ([28, 29, 30]).

Given the limited number of training images, larger architecture models would likely achieve even higher accuracy if trained with more samples. Overall, higher-capacity models appear capable of extracting background-invariant features even with limited data. Their performance seems constrained more by the amount of training data than by model capability; with additional images per individual, models such as `ViT-small` or `ConvNeXt-tiny` could reach near-perfect accuracy even under varying backgrounds.

Although some components, such as `Namer` and `sampleImgExtract.py`, offer user-friendly GUI interfaces, the current implementation requires a degree of technical expertise — namely familiarity with Python, 3D part design, and basic electronics integration with Raspberry Pi. However, these requirements are becoming increasingly manageable as AI-assisted coding tools and widely available modular maker technologies continue to lower the barrier for researchers without strong engineering backgrounds. We therefore remain optimistic about the system’s operability and broader adaptability.

## Acknowledgments

This work was supported by the Vienna Science and Technology Fund (WWTF) project ANIML (LS23-014).

## Appendices

### A Systemd service configuration; mAviary.service

~~~
[Unit]
Description=Aviary monitor
After=network.target
[Service]
Type=simple
User=brpi
WorkingDirectory=/home/brpi/code4Ethology/budgie
ExecStart=/bin/bash -c ‘source␣/home/brpi/mAviary/bin/activate␣&&␣python␣/
   home/brpi/code4Ethology/budgie/monitorAviary.py’
Restart=always
[Install]
WantedBy=multi-user.target
~~~

### B Confusion matrix of ConvNeXt-tiny model on metric_D2_b

|  | Afina | Famine | Fourier | Hades | Hilbert | Houdini | Lemon | Moonbeam | Pebbles | Storm | Summer |
| --- | --- | --- | --- | --- | --- | --- | --- | --- | --- | --- | --- |
| Afina | 29 | 0 | 0 | 0 | 0 | 0 | 0 | 0 | 0 | 0 | 1 |
| Famine | 0 | 30 | 0 | 0 | 0 | 0 | 0 | 0 | 0 | 0 | 0 |
| Fourier | 0 | 0 | 30 | 0 | 0 | 0 | 0 | 0 | 0 | 0 | 0 |
| Hades | 0 | 0 | 0 | 30 | 0 | 0 | 0 | 0 | 0 | 0 | 0 |
| Hilbert | 0 | 0 | 0 | 0 | 30 | 0 | 0 | 0 | 0 | 0 | 0 |
| Houdini | 0 | 0 | 0 | 0 | 0 | 30 | 0 | 0 | 0 | 0 | 0 |
| Lemon | 0 | 0 | 0 | 0 | 0 | 0 | 30 | 0 | 0 | 0 | 0 |
| Moonbeam | 0 | 0 | 0 | 0 | 0 | 0 | 0 | 29 | 1 | 0 | 0 |
| Pebbles | 0 | 0 | 0 | 0 | 0 | 0 | 0 | 0 | 30 | 0 | 0 |
| Storm | 0 | 0 | 0 | 0 | 0 | 0 | 0 | 0 | 0 | 30 | 0 |
| Summer | 0 | 0 | 0 | 0 | 0 | 0 | 0 | 0 | 0 | 0 | 30 |

## References

[1] Suresh Neethirajan. Recent advances in wearable sensors for animal health management. Sensing and Bio-Sensing Research, 12:15–29, 2017.

[2] Penny Hawkins, Sharon Brookes, James Bussell, Ngaire Dennison, Helmut Ehall, Anne-Marie Farmer, Theresa Langford, Chris Lelliott, Elliot Lilley, Ian Ragan, et al. Avoiding mortality in animal research and testing. Avoiding Mortality in Animal Research and Testing. University of Cambridge: RSPCA Research Animals Department, RSPCA Research Animals Department, 2019.

[3] Johnny P Do, Erwin B Defensor, Christine V Ichim, Maria A Lim, Jordan A Mechanic, Mark D Rabe, and Laura R Schaevitz. Automated and continuous monitoring of animal welfare through digital alerting. Comparative Medicine, 70(4):313–327, 2020.

[4] Kay E Earle and Nigel R Clarke. The nutrition of the budgerigar (melopsittacus undulatus). The Journal of nutrition, 121:S186–S192, 1991.

[5] EJ Burton, R Newnham, SJ Bailey, and LG Alexander. Evaluation of a fast, objective tool for assessing body condition of budgerigars (m elopsittacus undulatus). Journal of Animal Physiology and Animal Nutrition, 98(2):223–227, 2014.

[6] Luigi Fontana and Frank B Hu. Optimal body weight for health and longevity: bridging basic, clinical, and population research. Aging cell, 13(3):391–400, 2014.

[7] Zhuoyi Wang, Saeed Shadpour, Esther Chan, Vanessa Rotondo, Katharine M Wood, and Dan Tulpan. Asas-nanp symposium: Applications of machine learning for livestock body weight prediction from digital images. Journal of animal science, 99(2):skab022, 2021.

[8] M. Sharman Hoppes. Management of pet birds, 2024. In MSD Veterinary Manual, Merck & Co., Inc. (Originally published 2021, reviewed 2024).

[9] Andrew C Gallup, Elaine Herron, Janine Militello, Lexington Swartwood, Carmen Cortes, and Jose R Eguibar. Thermal imaging reveals sizable shifts in facial temperature surrounding yawning in budgerigars (melopsittacus undulatus). Temperature, 4(4):429–435, 2017.

[10] NR Augspurger and M Ellis. Weighing affects short-term feeding patterns of growing-finishing pigs. Canadian journal of animal science, 82(3):445–448, 2002.

[11] Temple Grandin and Chelsey Shivley. How farm animals react and perceive stressful situations such as handling, restraint, and transport. Animals, 5(4):1233–1251, 2015.

[12] Luca Catarinucci, Riccardo Colella, Luca Mainetti, Luigi Patrono, Stefano Pieretti, Ilaria Sergi, and Luciano Tarricone. Smart rfid antenna system for indoor tracking and behavior analysis of small animals in colony cages. IEEE Sensors Journal, 14(4):1198–1206, 2013.

[13] Natasha Dean Harrison and Ella L Kelly. Affordable rfid loggers for monitoring animal movement, activity, and behaviour. Plos one, 17(10):e0276388, 2022.

[14] Sabine G Gebhardt-Henrich, Alexander Kashev, Matthew B Petelle, and Michael J Toscano. Validation of a radio frequency identification system for tracking location of laying hens in a quasi-commercial aviary system. Peer Community Journal, 3, 2023.

[15] Roman Bumbálek, Jean de Dieu Marcel Ufitikirezi, Tomáš Zoubek, Sandra Nicole Umurungi, Radim Stehlík, Zbyněk Havelka, Radim Kuneš, and Petr Bartoš. Computer vision-based approaches to cattle identification: A comparative evaluation of body texture, qr code, and numerical labelling. Czech Journal of Animal Science, 70(9):383–396, 2025.

[16] YA Oldenhof. Effects of different marking techniques on the fitness of avian species. PhD thesis, Faculty of Science and Engineering, 2015.

[17] Lorenzo Pérez-Rodríguez, Roger Jovani, and Martin Stevens. Shape matters: animal colour patterns as signals of individual quality. Proceedings of the Royal Society B: Biological Sciences, 284(1849):20162446, 2017.

[18] Fernando Nottebohm. The origins of vocal learning. The American Naturalist, 106(947):116–140, 1972.

[19] Susan D Brown and Robert J Dooling. Perception of conspecific faces by budgerigars (melopsittacus undulatus): I. natural faces. Journal of Comparative Psychology, 106(3):203, 1992.

[20] Micheal L Dent and Robert J Dooling. Investigations of the precedence effect in budgerigars: The perceived location of auditory images. The Journal of the Acoustical Society of America, 113(4):2159–2169, 2003.

[21] Marisa Hoeschele and W Tecumseh Fitch. Phonological perception by birds: budgerigars can perceive lexical stress. Animal cognition, 19(3):643–654, 2016.

[22] Bernhard Wagner, Dan C Mann, Shahrzad Afroozeh, Gabriel Staubmann, and Marisa Hoeschele. Octave equivalence perception is not linked to vocal mimicry: budgerigars fail standardized operant tests for octave equivalence. Behaviour, 156(5–8):479–504, 2019.

[23] Alison Campbell and James W Tanaka. Inversion impairs expert budgerigar identity recognition: a face-like effect for a nonface object of expertise. Perception, 47(6):647–659, 2018.

[24] Rhea Diamond and Susan Carey. Why faces are and are not special: an effect of expertise. Journal of experimental psychology: general, 115(2):107, 1986.

[25] Keith M Kendrick, Khia Atkins, Michael R Hinton, Paul Heavens, and Barry Keverne. Are faces special for sheep? evidence from facial and object discrimination learning tests showing effects of inversion and social familiarity. Behavioural processes, 38(1):19–35, 1996.

[26] Angela Medina-García, Jodie M Jawor, and Timothy F Wright. Cognition, personality, and stress in budgerigars, melopsittacus undulatus. Behavioral Ecology, 28(6):1504–1516, 2017.

[27] Jinook Oh and Sylvia Cremer. ALTAA: analysis of long-term activity patterns in ant colonies. Methods in Ecology and Evolution, 2026.

[28] Florian Schroff, Dmitry Kalenichenko, and James Philbin. Facenet: A unified embedding for face recognition and clustering. In Proceedings of the IEEE conference on computer vision and pattern recognition, pages 815–823, 2015.

[29] Stefan Schneider, Graham W Taylor, and Stefan C Kremer. Similarity learning networks for animal individual re-identification-beyond the capabilities of a human observer. In Proceedings of the IEEE/CVF winter conference on applications of computer vision workshops, pages 44–52, 2020.

[30] Vincent Miele, Gaspard Dussert, Bruno Spataro, Simon Chamaillé-Jammes, Dominique Allainé, and Christophe Bonenfant. Revisiting animal photo-identification using deep metric learning and network analysis. Methods in Ecology and Evolution, 12(5):863–873, 2021.

